# Dimerization thermodynamics govern biological phase separation

**DOI:** 10.1101/2022.12.02.518839

**Authors:** J. Matthew Dubach

## Abstract

Many intrinsically disordered proteins/regions (IDR) can phase separate into a liquid state (biomolecular condensates) and/or a solid state (aggregates and amyloids). De-mixing in liquid-like phase separation carries an entropic cost that must be balanced by an energetic advantage of condensing. Yet, the mechanism of this energetic advantage remains unclear. Here, it is shown that an increase in dimerization binding interactions through liquid-like condensation can create overall energetic minimums when a system phase separates. Yet, the dimer binding interactions must be rapid explaining why IDRs are so prominent in phase separations. Numerous experimental examples that can only be described through dimer interactions support the dimer model presented here. The transition from condensates to aggregates and amyloids under certain conditions also appears to be driven by dimer abundance. Overall, the dimer model provides a novel theory that solves the thermodynamic mechanism of protein phase separation into biomolecular condensates, aggregates and amyloids.

## Introduction

Intrinsic disorder is abundant in the proteome^1^. While IDR function remains less characterized than structured proteins^2^, the ability of IDRs to phase separate within the cell^3-5^ has stoked interest into why phase separation arises and how it impacts cellular function^6,7^. Biomolecular condensates are liquid-like phase separations that flow, merge and dissolve^8^, rapidly exchange components with the surrounding environment^9^, are dependent on environmental conditions^10^, and can incorporate oligonucleotides^11^ or multiple other proteins^12^. Aggregates and amyloids are solid structures that often cannot be solubilized and are thought to arise from protein misfolding^13^. Intriguingly, numerous IDR proteins that aggregate in neurodegenerative disease have also been found to form biomolecule condensates under certain conditions^14-16^, suggesting that liquid and solid phase separation may have a common energetic origin.

Thermodynamically, phase separation produces a lower entropy of mixing, therefore, since separation is spontaneous, energetic advantages that overcome this entropic cost are required. The reversibility of biomolecular condensates presents a seemingly graspable thermodynamic phenomenon whereas the expectation that aggregation is driven by protein misfolding^13^ suggests more complex protein folding thermodynamics are involved. Initial considerations of biomolecular condensate thermodynamics modeled their formation as pure liquid-liquid phase separation (LLPS)^8^. But aspects of condensates do not fit canonical LLPS models^17,18^, therefore properties of polymer physics, such as Flory-Huggins interaction energies^19,20^, have been used to describe the energetic landscape of condensates. Discovery that the abundance of polar amino acids found in IDRs are drivers of condensate formation^21,22^ suggested that Flory-Huggins was a good model for condensation. Yet, observations that altered order of amino acids^23^ and replacement of charged amino acids^24^ prevents condensate formation led to the “stickers and spacers” model where LLPS, multivalent interactions^25^, and percolation govern condensation^21,26^. Sequence dependency on dense phase concentration and material properties of condensates^27,28^ as well as loss of just a single sticker impacting the concentration at which condensates arise^29^ suggest that binding interactions are essential. However, in the sticker and spacer model, why condensates spontaneously form, the impact of the complex biological environment^18^ and why single amino acid mutations^24^ disrupt condensate formation remain unanswered. Furthermore, how IDRs can form both condensates and amyloids remains largely unexplained^30^. Tau amyloids can be highly structured^31^, suggesting that binding interactions in solid phase separation can be very specific and not complex entanglements.

Recent results show that IDR proteins can bind to form dimer pairs with high affinity and rapid dynamics^32-34^. However, the rapid dynamics of IDR protein interactions prevent *de novo* discovery in typical protein interaction pulldown screens. Thus, IDR binding interactions need to be explicitly measured, which likely limits recognition of the prevalence of these interactions in biology. Therefore, here, the energetic impact of dimerization is considered as a driver of phase separation (the dimer model).

## Results

### Dimerization thermodynamics predict observed phase separation

Similar to disordered proteins and peptides, low molecular weight compounds can undergo liquid-like phase separation^35^. One molecule, diphenylalanine capped with a methoxy group (FFome), undergoes phase separation with a pH dependency^36^. At low pH the amine is protonated and the dipeptide is soluble up to 50 mM, remaining diffuse (Fig. S1A). At 25 mM FFome is diffuse at pH 3.5, forms liquid-like phase separation at pH 9 and forms aggregate crystals at pH 13 (**Fig. 1A**). The formation of condensates at pH 9 is also dependent on concentration with a decreasing abundance at 10 mM, and, surprisingly, less abundance at 45 mM (**Fig. 1B**). FFome phase separation is very dynamic as well. A rapid and large increase in pH transitions the system from diffuse to formation of condensates to crystal structures within seconds (Fig. S1B). These dynamics are highlighted by local addition of base into a diffuse solution (**Fig. 1C**). Locally raising pH by adding 2 μl of 10 M NaOH to 100 μl 50 mM FFome at pH 2 generates condensates at the site of addition, yet over time condensates dissolve as the NaOH diffuses away and mixes with the rest of the solution, producing a solution pH < 3. However, some crystal aggregates are formed and remain in solution demonstrating that transient deprotonated FFome formed a small fraction of irreversible structures similar to those observed at stable high pH.

**Figure 1.**
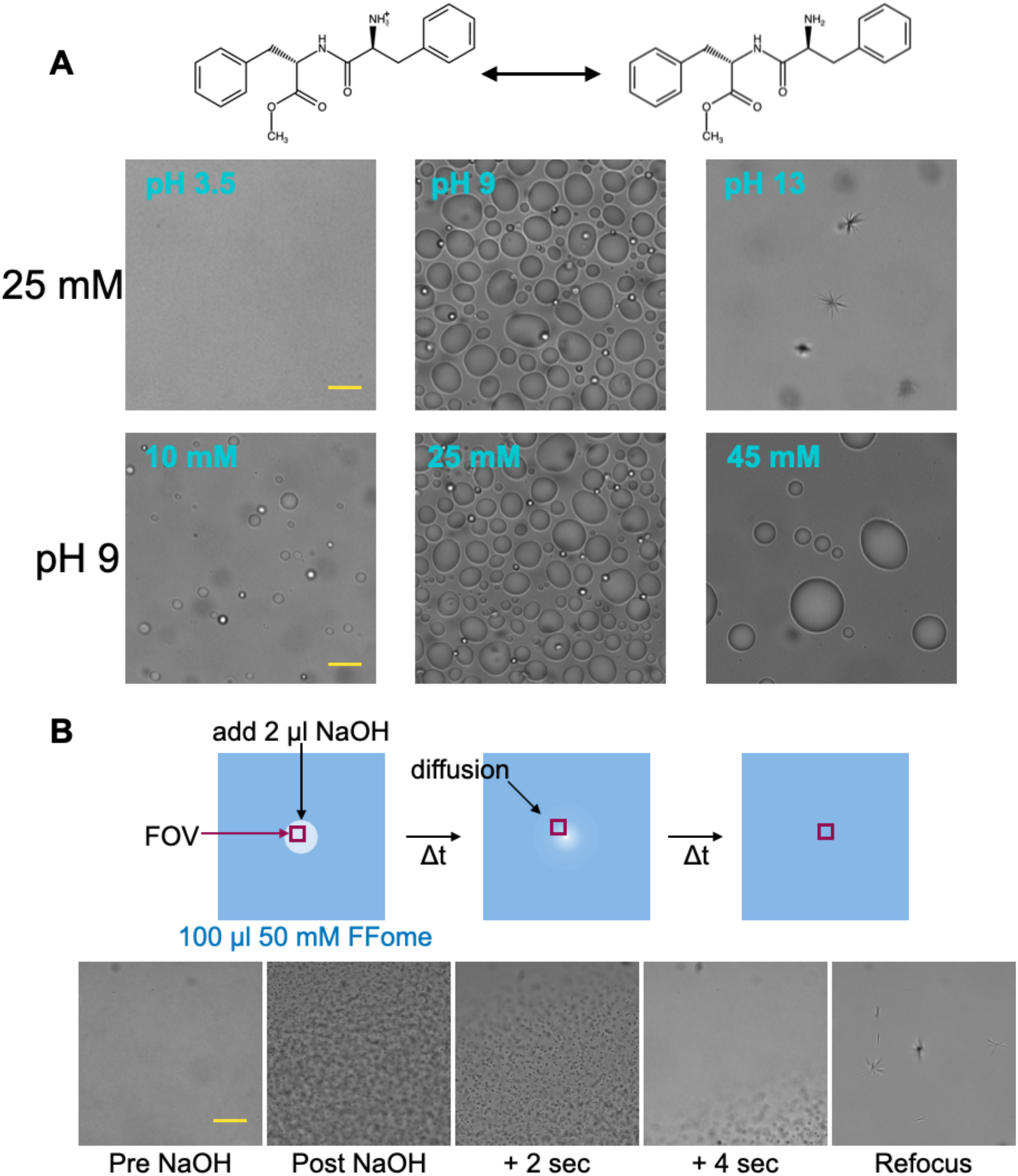
Phase separation of FFome. (**A**) FFome will be protonated in a pH-dependent manner. Representative DIC images of FFome in solution at specific pH and concentrations, scale bars = 30 μm. (**B**) Schematic for local addition of high concentration NaOH in a solution of FFome. Representative DIC images of FFome before NaOH addition and post addition and multiple time points as well as refocusing in the sample after condensation dissolved, scale bar = 40 μm.

Why FFome phase separates in a pH dependent manner is unknown. The small size of the dipeptide suggests that simple binding interactions must be drivers of phase separation. Since all molecules have an intermolecular force of interacting^37^, FFome in solution has a binding affinity to form a homo-dimer, which can be expressed as a dissociation constant (K_d_). Thermodynamically, the K_d_ is a measure of the decrease in Gibbs free energy of the system that arises through binding. The steady-state fraction bound for any interacting molecules can be plotted as a function of the total concentration to K_d_ ratio ([protein]/K_d_), where higher ratios generate more binding (**Fig. 2A**). Critically, since the bound fraction is a steady-state value it applies to interactions with any binding dynamics. It is expected that protonated, positively charged FFome has a lower dimer binding affinity than deprotonated FFome. Likewise, the abundance of FFome impacts how much exists in a transient dimer state. Therefore, FFome concentration and protonation state, two aspects that govern condensation experimentally, will impact the dimerization fraction as well.

**Figure 2.**
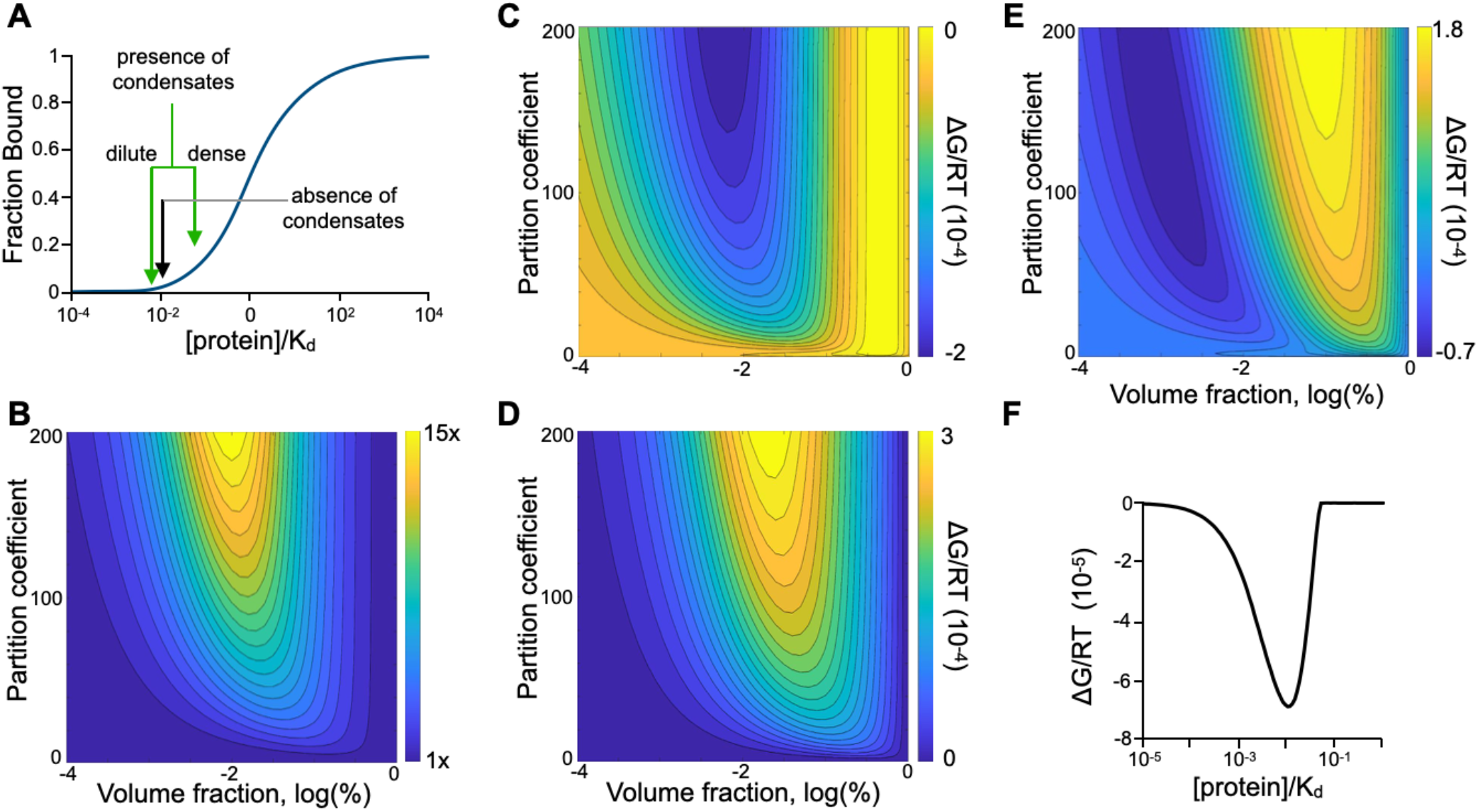
The dimer model of phase separation predicts system energy minimums upon liquid-like condensation within a limited [protein]/K_d_ ratio range. (**A**) The steady state fraction bound as a function of the [protein]/K_d_ ratio. Condensation generates two phases that reside on the fraction bound curve. (**B**) Calculations for total homo-dimerization fraction bound upon liquid-like phase separation as a function of dense phase volume fraction and partition coefficient. (**C**) Calculations for the change in the Gibbs free energy of binding upon liquid-like phase separation as a function of dense phase volume fraction and partition coefficient. (**D**) Calculations for the change in the entropy of de-mixing upon liquid-like phase separation as a function of dense phase volume fraction and partition coefficient. (**E**) Calculations for the change in total system energy upon liquid-like phase separation as a function of dense phase volume fraction and partition coefficient. (**F**) Calculations for the minimum change in total energy that can be achieved through liquid-like phase separation as a function of the [protein]/K_d_ ratio of homo-dimerization. B-F, K_d_ = 10 mM.

Liquid-like phase separation generates a high concentration dense phase and a low concentration dilute phase, which is described by the partition coefficient ([dense]/[dilute]). Through a higher concentration the dense phase will have a higher abundance of dimerization than the dilute phase (**Fig. 2A)**. Mass balance calculations (supplemental information) of total steady-state fraction bound as dimers in a system at a [protein]/K_d_ ratio of 0.01 reveals that phase separation produces an overall higher fraction bound (**Fig. 2B**) - the amount of binding is dependent on the partition coefficient and volume fraction of the dense phase. Increased dimer abundance in the presence of condensates generates a lower effective dissociation constant, which can be used to calculate the impact on the Gibbs free energy of binding (**Fig. 2C &** supplemental information). Here, the system has a boundary condition that limits the total concentration possible in the dense phase (supplementary information).

However, the entropic cost of phase separation generates an increase in system free energy that is also dependent on the volume fraction and partition coefficient (**Fig. 2D &** supplemental information). Yet, this entropic cost is shifted from the Gibbs free energy of binding in partition coefficient vs. volume fraction space. Summing the two energies reveals that liquid-like phase separation in system with a [protein]/K_d_ ratio of 0.01 creates an energetic minimum state (**Fig. 2E**). Here, the energetic advantage of phase separation achieved by increased dimer abundance overcomes the entropic cost of de-mixing at specific partition coefficients and volume fractions. Calculating the minimum achievable energy through condensation as a function of [protein]/K_d_ ratio demonstrates that phase separation is only energetically favorable within a specific range of [protein]/K_d_ ratios (**Fig. 2F**). At low and high [protein]/K_d_ ratios the reduction in energy through increased total dimer abundance upon phase separation does not overcome the entropic cost at any volume fraction and partition coefficient. This limited range suggests that 25 mM FFome at pH 3.5 is below the phase separating [protein]/K_d_ ratio, while 25 mM FFome at pH 13 is above the liquid-like phase separating ratio. Overall, these calculations suggest that condensation can be driven by simple dimer binding interactions.

### Hetero-interacting peptides behave in a manner predicted by the dimer model

Hetero-interacting molecules can also undergo phase separation. Using previously demonstrated^38^ simple mixtures of polylysine (polyK) and polyglutamic acid (polyE), phase separation was confirmed to be dependent on the peptide length (**Fig. 3A**). Equimolar mixtures of 50mer peptides formed aggregates at the concentrations where 30mer and 20mer peptides form condensates, while 4mer peptides showed no phase separation. The 20mer and 30mer peptide condensates exhibited both merging and spontaneous disappearance (Fig. S4A). To determine if aggregation of the 50mer peptides arose from excess mass, lower concentrations of equimolar mixtures of 50mer polyK and polyE were generated, but these also produced aggregates (**Fig. 3B**). However, mixtures of equimolar E_30_ and E_20_ with K_50_ formed condensates at 2 mM (**Fig. 3C**). Thus, despite the same overall abundance of lysine and glutamic acid as the 50mer mixture at 2 mM, simply cutting one peptide into two sections transitioned the mixture from solid aggregates to liquid-like condensates. Charge neutral mixtures of 20mer or 30mer polyE with K_50_ also formed condensates at 2 mM K_50_ (**Fig. 3C**). Yet, charge neutral mixtures of E_4_ with K_50_ failed to form any phase separation at 20 mM K_50_ (Fig. S4B). Aggregates and not condensates formed at higher concentrations of equimolar E_30_, E_20_ and K_50_ mixtures (**Fig. 3C**). Aggregates also formed at higher concentrations of charge neutral mixtures of E_30_ and K_50_ and E_20_ and K_50_ (**Fig. 3C**).

**Figure 3.**
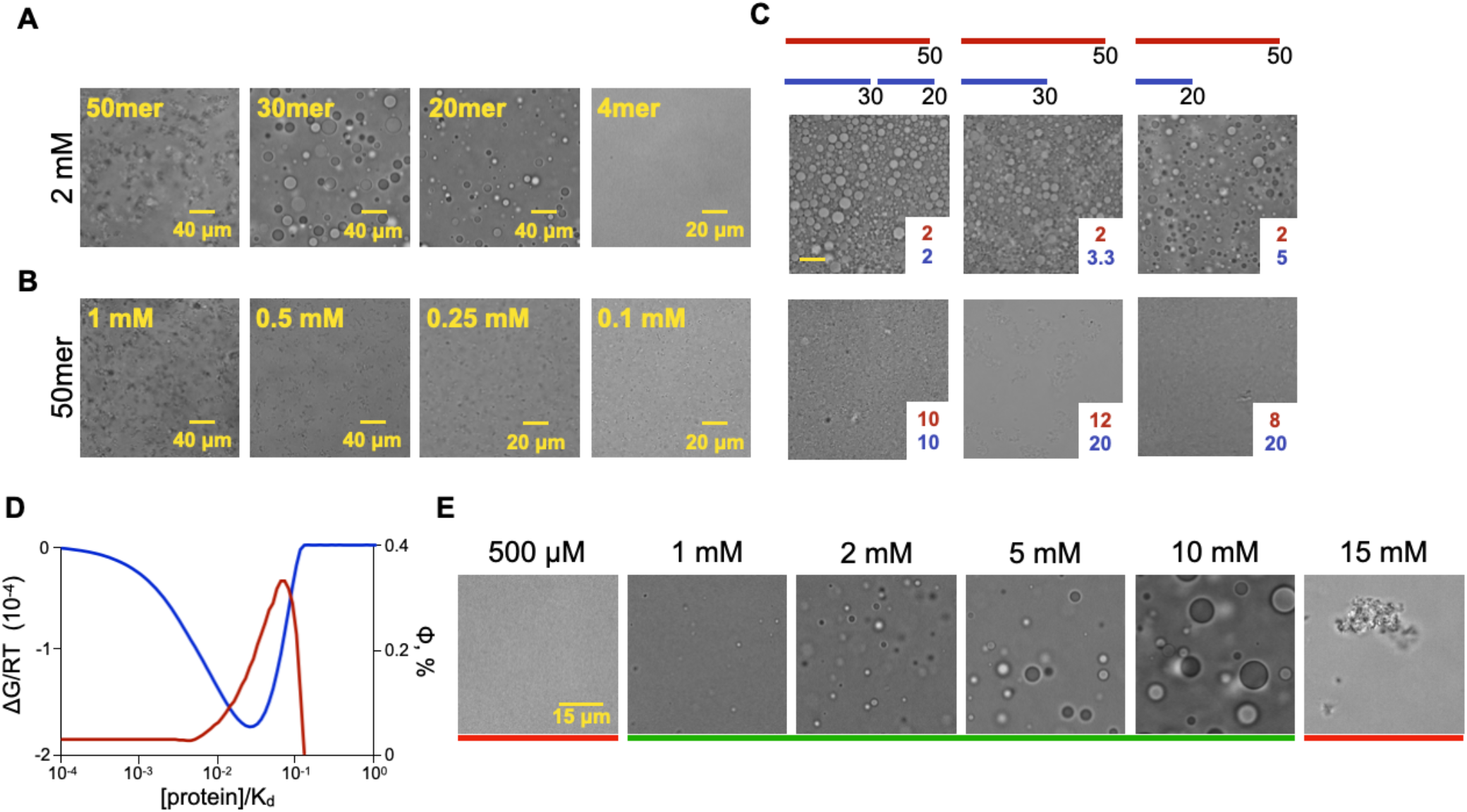
Polyion condensate formation follows the dimer model predictions. (**A**) Representative DIC images of equimolar polyK and polyE mixtures at 2 mM. (**B**) Representative DIC images of equimolar K_50_ and E_50_ mixtures at multiple concentrations. (**C**) Representative DIC images K_50_ (red) and polyE (blue) mixtures at equal charge ratios, shown overlaying each image is the concentration of K_50_ (red) and polyE (blue) in mM. (**D**) Calculated energy minimum for condensate formation (blue) and volume fraction (red) at the minimum energy as a function of protein concentration (single protein) for an equimolar mixture of two binding proteins, K_d_ = 10 mM. **(E)** Representative DIC images of K_20_-E_20_ peptide condensates at equimolar concentrations in the presence of 350 mM NaCl, with red and green indicating the absence or presence of condensates, respectively.

At charge neutrality the only difference between E_30_ and/or E_20_ mixed with K_50_ compared to E_50_ mixed with K_50_ is the length of the polyE, suggesting that length and not capacity to form multivalent interactions is the critical factor dictating phase separation. PolyK and polyE in solution can bind to form dimers with binding affinities dependent on the length of the peptide^39^, thus 50mer peptides will dimerize with a higher affinity than 30mer or 20mer peptides. Therefore, mixtures of E_30_ and E_20_ with K_50_ will have a lower dimer abundance than mixtures of E_50_ with K_50_. The condensation dependency on both length and concentration suggests that the dimer abundance of peptide mixtures governs the ability to form liquid-like condensates in the same manner as FFome interactions.

Similar to homo-binding interactions, energy calculations of hetero-interactions predict a [protein]/K_d_ ratio range where condensates are energetically favorable (**Fig. 3D**, Fig. S5 & supplemental information). Furthermore, the predicted volume fractions that achieve the energetic minimum are calculated to remain below 1%. These findings are in stark contrast to existing theories that predict condensates will increase in size with increasing concentration until the dense phase becomes system spanning. Prior research on the concentration dependence of condensate formation commonly describes the concentration above which condensates appear (C_sat_)^24^ but does not establish a system spanning dense phase. Fortunately, the concentration dependence of condensate dissolution, as predicted here in the dimer model, is readily testable through increasing the protein concentration of a system with stable condensates. Thus, equimolar mixtures of K_20_ and E_20_ peptides in 350 mM NaCl were created at increasing peptide concentrations. In this system, condensation demonstrated a C_sat_ at 1 mM of each peptide, but also dissolution at 15 mM of each peptide (**Fig. 3E**), before system spanning dense phase predicted by the existing model.

### Polyion peptide phase separation is driven by dimerization

Salt can induce condensate reentrance for phase separating peptides with charged amino acids^40^, suggesting that the abundance of ions impacts the energy of condensate formation. Intriguingly, dimerization of two oppositely charged polyions primarily arises from the gain in entropy of counterion release and can be disrupted by high counterion concentrations that lower the energetic advantage of binding^39^. Therefore, if binding is the driver of condensate formation, the presence of counterions will disrupt the energetic advantage of condensation. To test if counterion concentration impacts condensate formation, equimolar mixtures of K_20_ and E_20_ peptides at various peptide and salt concentrations were created and the presence of condensation was determined through DIC imaging. An automated image analysis algorithm quantifying the non-background pixels was used to identify the presence of condensates (Fig. S4C). A plot of condensation as a function of salt and peptide concentration demonstrates that condensates reenter solution with a similar energy pattern to the molar free energy calculated from the dimer model for hetero-dimer interactions (**Fig. 4A & 3D**). At higher concentrations, condensates formed upon vigorous mixing but lost homogenous condensation within 10 minutes (Fig. S6A-D). The volume fraction was also dependent on both the peptide concentration and salt concentration (**Fig. 4B**), as predicted by the dimer model. The formation of condensates as a function of salt and peptide concentration was also determined for K_30_ and E_30_ at equimolar ratios. The 30mer reentrance boundary curve (**Fig. 4C**) displayed a shift toward lower peptide concentrations and higher salt concentrations compared to the 20mer curve, suggesting a higher dimer affinity as expected. Similar results were found with binding between 1:1 mixtures of K_10_ and D_10_, where condensates were disrupted by lower salt concentrations (**Fig. 4C**). However, for the 10mer peptides, a sharp decrease in the reentrance boundary curve at high concentrations could not be established due to limiting counterion concentrations in the peptide stock solutions. Yet, 10mer peptide condensates at the upper concentration limit generated smaller condensates (Fig. S7E), similar to observed salt concentration reentrance of 20mer peptides (**Fig. 4B**), confirming the impact of counterions at these concentrations. Critically, the correlation of peptide length with salt concentration needed to prevent condensate formation confirms that phase separation proceeds through dimer interactions. Otherwise, if phase separation arose through multivalent interactions, for example, the concentration of salt that prevents condensation at 1 mM 30mer peptides would be the same as the concentration that disrupts 1.5 mM 20mer peptides (equal total monomer concentration), but that is not the case.

**Figure 4.**
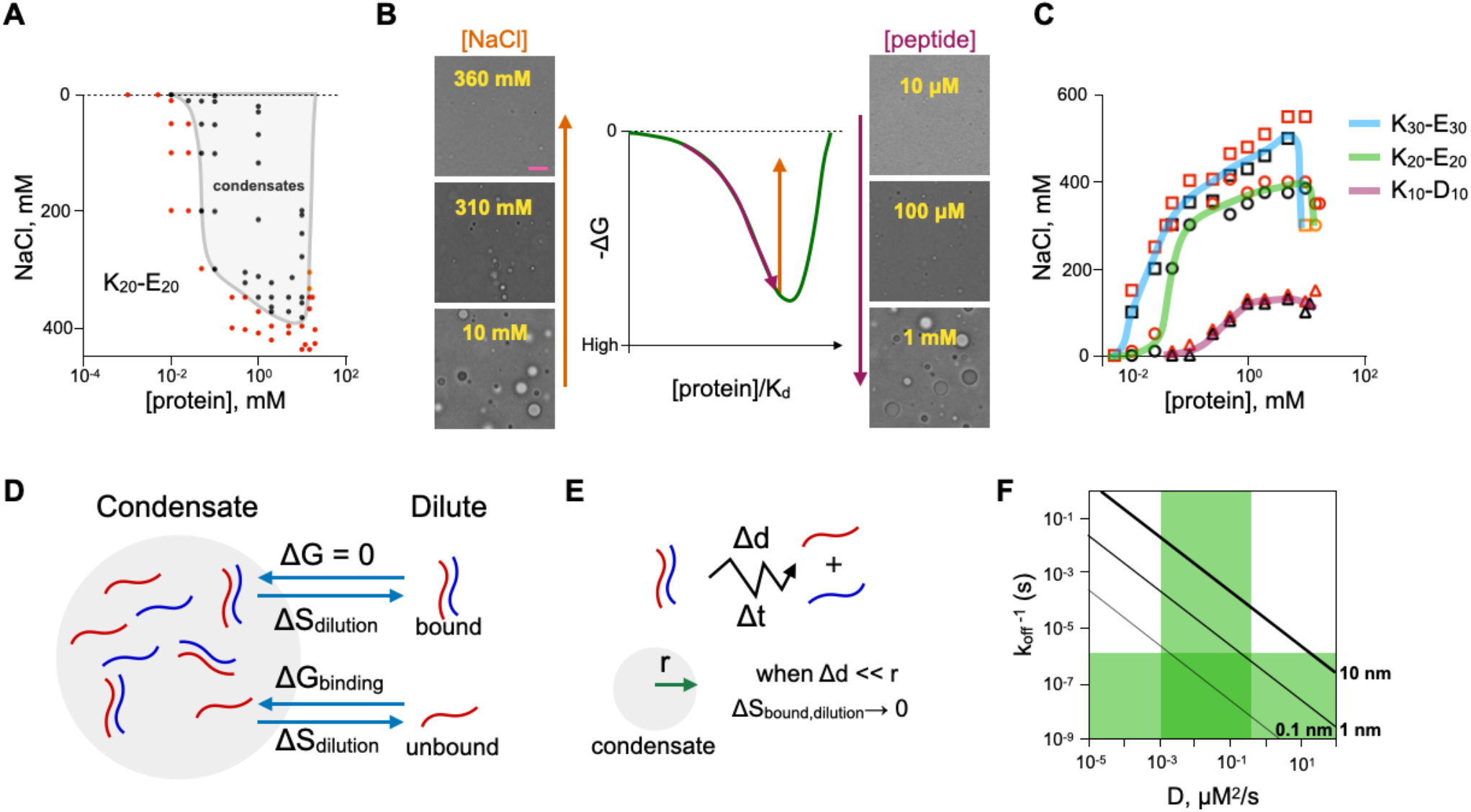
Condensate salt disruption and binding dynamics in the dimer model. (**A**) Formation (black) or absence (red) of K_20_-E_20_ condensates at varying NaCl concentrations and equimolar peptide concentrations. A trace between the presence and absence of condensates highlights condensate forming conditions. Orange data points are condensates that stabilize into aggregates. (**B**) Representative DIC images of K_20_-E_20_ peptide condensates as a function of NaCl (orange, 1 mM peptide) and peptide (purple, [NaCl] = 20x[peptide]) concentration, scale bar = 10 μm. (**C**) Presence (black) and absence (red) of condensates for K_20_-E_20_ (circles – green trace), K_30_-E_30_ (squares – blue trace) and K_10_-D_10_ (triangles – purple trace) with traces of the boundary for condensate reentrance as a function of [NaCl] and equimolar peptide concentration. (**D**) The energetic forces impacting bound and unbound molecules that form condensates. **(E)** The dissociation rate and diffusivity govern the diffusion distance of a bound molecule. (**F**) Reported (green) dissociation rates of intrinsically disordered proteins in condensates and diffusivity measured in condensates. Isolines (black), determined by Fick’s law of diffusion, are plotted for average diffusion distance of bound protein before dissociation.

The mechanism of binding-induced counterion release driving condensate formation was further confirmed in unequal molar ratios of E_30_ and K_30_ (Fig. S7A). Here, counterions from the stock solution were unbalanced and condensates were not formed. However, addition of NaCl, which provided ions for release upon binding, enabled condensate formation at these unbalanced ratios. The dimer model also predicts that the presence of one condensate component above the dissolution concentration will prevent condensate formation (Fig. S7B). This was confirmed experimentally with K_20_ and E_20_ where E_20_ was held above the dissolution concentration and lower concentrations of K_20_ were unable to generate condensates (Fig. S7C). However, when both E_20_ and K_20_ were at concentrations below the dissolution concentration, the imbalanced ratio from lowering the concentration of just K_20_ did not prevent condensate formation (Fig. S7C).

### Binding affinity and dynamics govern condensation and aggregation

Seemingly, both the dimer model presented here and the existing theory of biomolecular condensates do not provide compelling mechanisms for aggregate and not condensate formation observed with 50mer peptides. Decreasing the peptide concentration reduced the size of the aggregates but did not generate condensates (**Fig. 3B**). Similarly, increasing the salt concentration also reduced the size of the aggregates, suggesting that aggregates form through polyion binding, but did not generate any condensates (Fig. S8). The NaCl concentration (850 mM) needed to prevent aggregates was higher than the concentration (500 mM) that prevented of E_30_-K_30_ condensates at the same peptide concentration, confirming higher affinity binding between the 50mer peptides.

Here, the kinetics of binding interactions that drive condensate formation in the dimer model are considered. Proteins readily exchange between the dense and dilute phase^41-47^, yet condensation is stable in steady state demonstrating that total flux is balanced. In the dimer model, condensates consist of both bound and unbound protein. In the binding model, unbound protein flux is balanced by the entropic force to leave the condensate and the energetic advantage of existing at a high condensate concentration to increase the likelihood of binding (**Fig. 4D**). However, bound protein does not have any energetic advantage to enter nor remain within the condensate and is only subject to the gain in entropy upon leaving, as well as a decrease in chemical potential at high dense phase concentrations. Since flux between the two phases is diffusion driven and the rates of diffusion for bound and unbound protein will be similar, this imbalanced force will seemingly prevent condensation. Yet, diffusion out of condensates is a dynamic process, and, over time, a bound protein both diffuses and dissociates (**Fig. 4E**). Recent results demonstrate that IDR protein binding in a condensate occurs through extremely fast dynamics, with partner exchange rates suggesting k_off_ rates greater than 10^6^ 1/s^48^. Measured diffusivity rates inside condensates have been reported between 10^−3^ to 10^−1^ μm^2^/s^41-46^. Therefore, it is expected that molecules within condensates can exist with dynamics where bound protein dissociates before significant diffusion relative to the size of the typical condensate (**Fig. 4F**). This phenomenon enables steady state condensate formation and suggests that the dissociation rate of K_50_-E_50_ is too slow to form condensates. Since both 50mer and shorter peptides are expected to have the same diffusion-limited association rates, the higher affinity of the longer peptides arises solely through a slower dissociation rate. Therefore, the slower dissociation rate prevents liquid-like condensate formation. Instead, dimerized 50mer polyK and polyE will be hydrophobic as the charged components are interacting with each other, thus aggregation likely forms when multiple dimers interact.

### The dimer model captures protein-oligonucleotide phase separation

Condensates can also be generated from mixtures of proteins and oligonucleotides^49,50^. Mixtures of K_20_ and a 20mer single stranded DNA (DNA_20_) produced condensates (**Fig. 5A**). The solubility limitations of DNA prevent 1:1 charge ratio at higher concentrations, thus determination of concentration induced condensate dissolution was performed at uneven molar mixtures. Dissolution was not observed at lower concentrations with the same charge ratio (**Fig. 5B**), confirming that loss of condensates at high concentration is driven by concentration and not charge ratio. Both the observed C_sat_ and dissolution concentration where higher for K_20_-DNA_20_ than K_20_-E_20_, suggesting that K_20_ binds DNA_20_ with a lower affinity than E_20_. However, mixtures of K_20_ and a 20mer RNA (RNA_20_) were unable to generate condensates, instead producing aggregates (**Fig. 5C**). Absence of condensates was also found for 1:1 charge ratios of RNA_20_ with K_50_, K_30_, K_10_ and K_4_ (**Fig. 5C**), with all except K_4_ producing aggregates.

**Figure 5.**
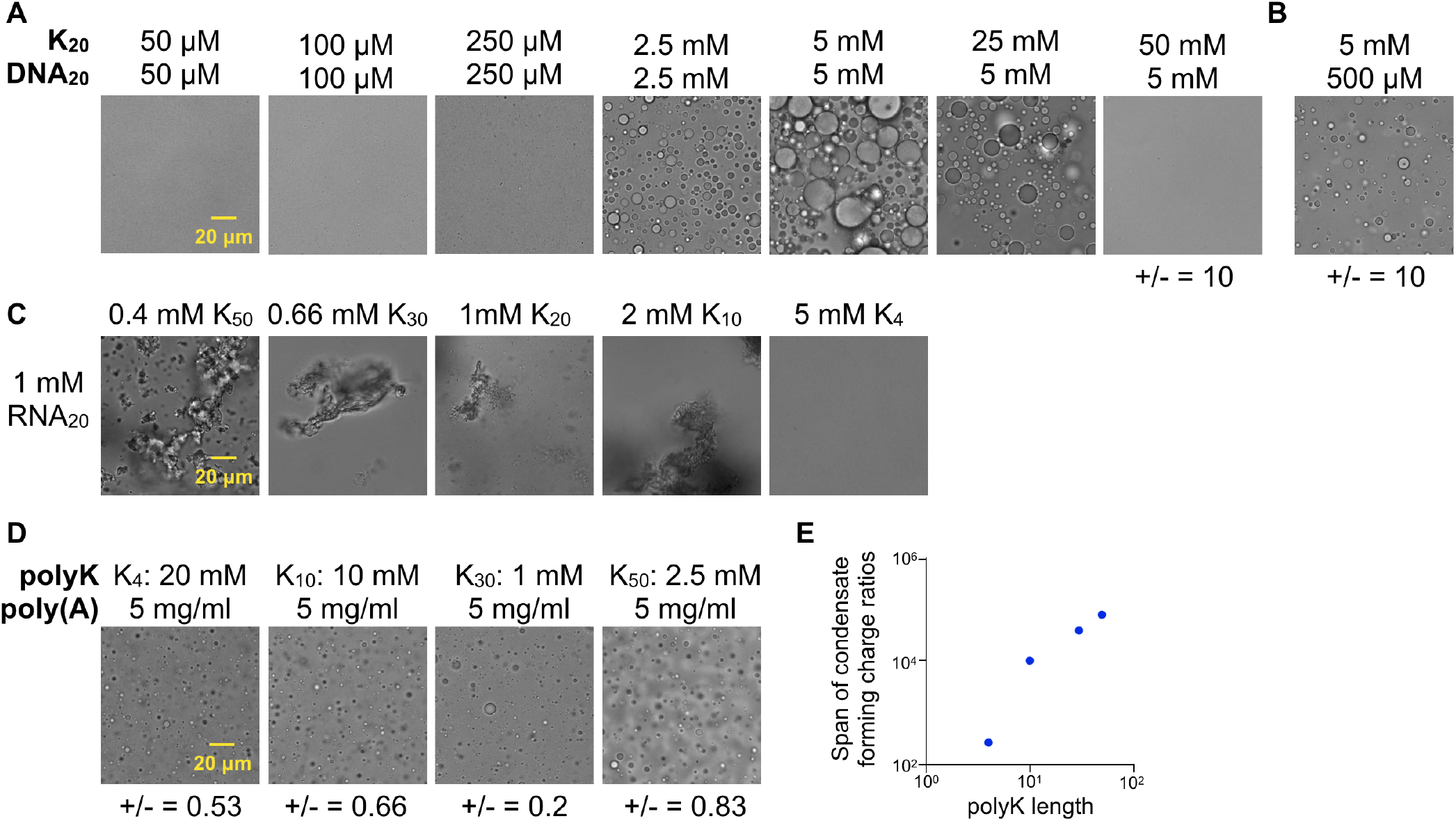
Condensates formed by peptides and oligonucleotides follow the dimer model. (**A**) Representative DIC images of K_20_-DNA_20_ mixtures. (**B**) Representative DIC image of a K_20_-DNA_20_ mixture at equal charge ratio as the highest concentration in (A). (**C**). Representative DIC images of polyK-RNA_20_ mixtures. (**D**) Representative DIC images of polyK and poly(A) mixtures. (**E**) The charge ratio span (fold difference between the positive and negative extremes) for polyK and poly(A) mixtures over which condensates form as a function of polyK peptide length.

The absence of condensates remained at lower concentrations of polyK and RNA as well (Fig. S9), suggesting that polyK binding to RNA has a much higher affinity than binding to DNA or polyE.

Condensates were formed, however, between polyK and poly(A) RNA, a long RNA polymer, (**Fig. 5D**), demonstrating that polyK can form condensates with RNA. As a large polymer, a single molecule of poly(A) can bind multiple molecules of polyK. Therefore, each poly(A) molecule is unlikely to be saturated with polyK, leaving sections unbound, creating a more balanced force to remain in the condensate (**Fig. 4D**). The dimer model also predicts that for binding interactions with one molecule having multiple binding sites the dissolution concentration is higher than for one-to-one binding (Fig. S10A). This increased dissolution concentration arises from the lower entropic cost of RNA condensation. All polyK peptides showed loss of condensation at both high and low polyK:poly(A) concentration ratios (Fig. S10B). Existing models of condensate formation between oppositely charged multivalent molecules have predicted that charge imbalance is the driver of dissolution^51-55^. However, intriguingly, here the span of charge ratios where condensates were stable was dependent on the length of the polyK peptide (**Fig. 5E**), demonstrating that reentrance with multimeric interactions is likely due to dimer forming thresholds and not a charge imbalance. At extreme concentration imbalances the likelihood of the lower concentration component being bound in solution is much higher than at equal ratios. Therefore, the energetic advantage of phase separation versus diffuse distribution is lost at large concentration imbalances and the system remains diffuse.

### Polyions can form amyloid fibrils at high dimer abundance

Experimental results with polyion peptides demonstrate that high concentrations where condensates are not present generated aggregates. For the 50mer peptides, which do not form condensates at any concentration tested, aggregation produced higher ordered structure with increasing concentration (**Fig. 6A**). And, while difficult to quantify, higher order structures appeared to be more abundant at higher concentrations for 10mer, 20mer and 30mer polyion mixtures (**Fig. 6A**). Generation of these highly ordered, amyloid-like structures at high concentrations was immediate. However, concentrations close to the condensate dissolution concentration initially formed condensates upon peptide mixing that became smaller in size over time and eventually disappeared leaving small aggregates (**Fig. 6B**). This observed reduction in condensate size over time was similar to concentration dependent condensate size (**Fig. 3E**), suggesting that the soluble fraction of peptide was reduced over time as peptide formed aggregates.

**Figure 6.**
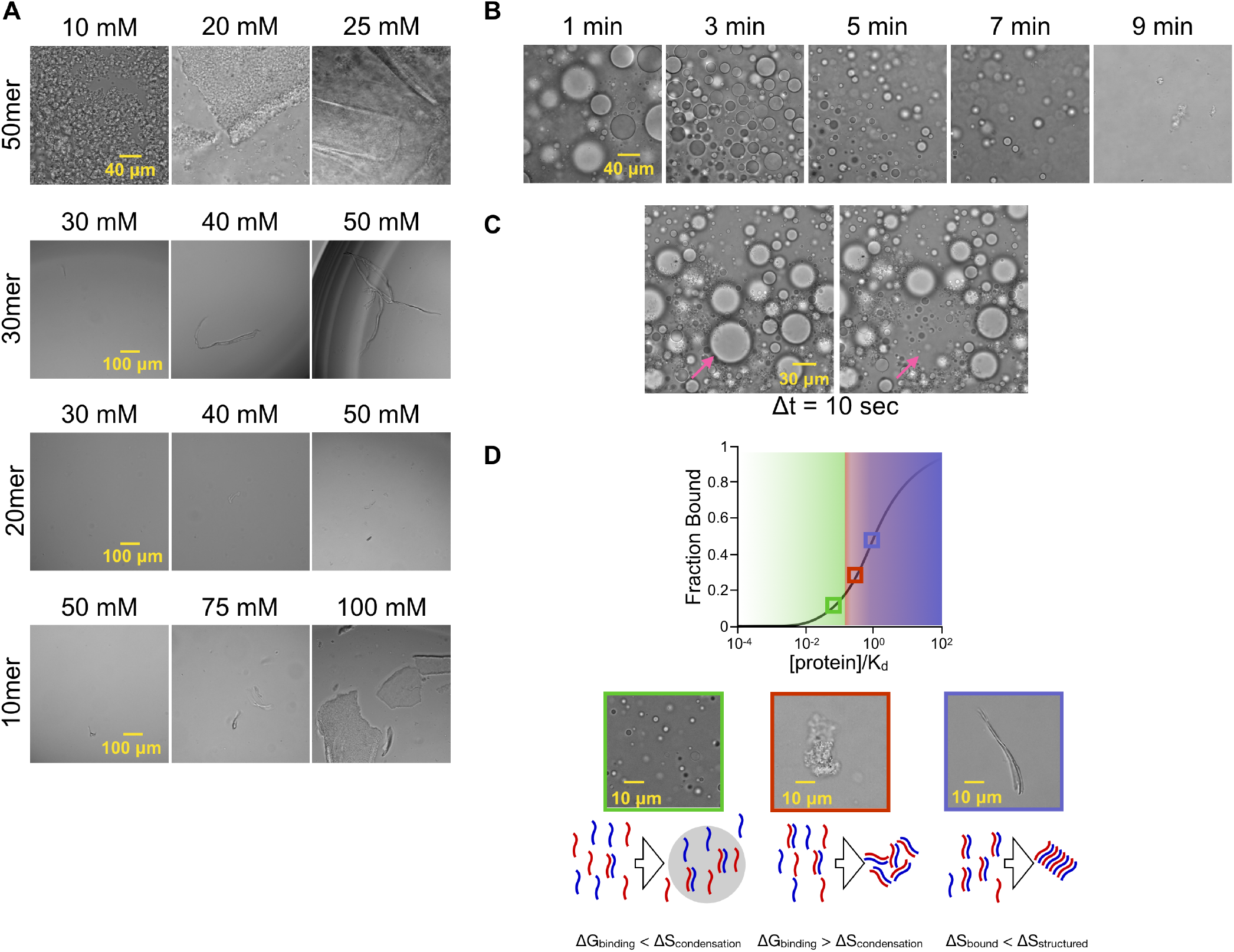
Polyions form structured aggregates at higher concentrations. (**A**) Representative DIC images of K_50_-E_50_, K_30_-E_30_, K_20_-E_20_, and K_10_-D_10_ equimolar mixtures. (**B**) Representative DIC images of K_20_-E_20_ equimolar mixture at 15 mM over time. (**C**) DIC images of K_30_-E_30_ equimolar mixture at 10 mM at 5 minutes and 10 seconds later, arrow indicating a condensate that dissolves. (**D**) Model of phase separation. At [protein]/K_d_ < 10^−1^, condensation produces an energy minimum through increased binding. Just above [protein]/K_d_ = 10^−1^, condensation is not energetically favorable and the steady state abundance of dimerized protein induces aggregation. At higher [protein]/K_d_ values more dimerized protein generates higher order structures such as amyloid fibrils.

Furthermore, dissolution of condensates during transition into aggregates did not directly produce aggregates but instead reentered solution (**Fig. 6C**), also suggesting that these condensates are not precursors to aggregates.

The dimer model and supporting experimental results suggest that the fraction of steady-state dimerized IDR governed by the [protein]/K_d_ ratio dictates phase separation (**Fig. 6D**). At low bound fractions the decrease in the free energy of binding that can be achieved through condensation overcomes the entropic cost of de-mixing and biomolecular condensates produce the lowest energetic state, provided the binding interactions are fast. However, at higher [protein]/K_d_ ratios the diffuse dimer abundance is also higher and the relative increase in binding within higher concentration condensates does not outweigh the entropic cost of de-mixing so condensates are not energetically favorable. Yet, the abundance of steady-state dimerized protein in solution enables a new energetic minimum when the dimerized IDR forms aggregates. Aggregation likely arises from the low entropy of solvent interacting with dimerized peptide outweighing the entropic cost of de-mixing. At even higher [protein]/K_d_ ratios, the increased abundance of dimerized IDR appears to enable a lower energetic minimum to be reached through more structured amyloid formation. Critically, with the peptides used here, aggregation is not a product of condensation, rather it occurs when condensates are not energetically favorable or dynamically possible, demonstrated by aggregation of the 50mer peptides (**Fig. 3A**).

## Discussion

The energetic advantage of dimerization, either homotypic or heterotypic has not been previously explored as a contributor to phase separation. However, binding interactions between highly flexible molecules are essential to condensate formation^56^. And, experimentally, condensation has been shown to be dependent on system temperature^57^, IDR phosphorylation^58^, and buffering of electrostatic interactions^3,40,59^, diverse properties that alter binding. The recent demonstrations that IDRs dimerize with extremely fast dynamics while maintaining a disordered state^32,33,48,60^ suggests that dimerization binding interactions are more abundant than currently appreciated. Here, consideration of the energetic impact of dimerization captures the majority of literature results on liquid-like condensates. The dimer model accurately captures salt induced reentrance^40^ and indicates that binding energy limitations at imbalanced molar ratios, and not charge imbalance^52^, induces dissolution, a finding that may clarify transcriptional condensate function^61^. The model also describes complex coacervation^62^ and the energetic forces governing condensation between oppositely charged molecules^51^. The reliance of condensate formation on very fast dissociation kinetics also explains why biomolecular condensates are nearly unique to disordered proteins.

The dimer model suggests that rather than physical crosslinks among multivalent proteins being the cause of condensate formation^63^, simple dimerization induces condensates. Therefore, sequence mutations in disordered proteins that prevent condensate formation^24^ do not remove stickers, but rather alter the binding affinity. However, the dimer model does not exclude the presence of multivalent interactions, when possible, within a condensate. The dimer model simply demonstrates that the energy gained from being bound to one or many molecules and the higher likelihood of being bound within a higher concentration environment drives phase separation. Indeed, multiple binding interactions that molecular dynamic simulations suggest can happen within a condensate^48^ could prevent aggregation.

Dimerization as a driver of condensation also demonstrates how IDRs can form aggregates and amyloids. Stacked β-sheets^64^ are the core component of fibrillar structures^31,65^, demonstrating that IDRs interact with precise alignment. Thus, the steady-state abundance of dimers at high [protein]/K_d_ ratios provides aligned precursor for amyloid stacking. And, the presence of aggregate seeds^66^ provides a substrate for dimers to grow fibrils through dimer-seed interactions at a rate that will be dependent on the [protein]/K_d_ ratio in the system. The dimer model, however, does not require that the entire length of an IDR be bound, congruent with the observed fuzziness of tau fibrils^67^.

Experimental measurements based on dimer model predictions also confirm that the dimer model accurately captures phase separation. Increasing protein or peptide concentrations well beyond C_sat_ has not been previously explored since existing models suggest condensates would persist, however the dimer model suggests otherwise and performing experiments at high peptide concentrations confirms this prediction. Several outstanding questions remain and the dimer model is certainly incomplete. The size of the condensing components will impact the condensate volume fraction, while the surface tension will impact the number and size of condensates. However, overall, the dimer model presented and validated here predicts observed phase separation that cannot be described by other theories.

Viewing IDR phase separation phenomena through the lens of the dimer model suggests that protein sequence^68^ and post translational modifications^58,69^ impact phase separation by altering binding affinities – a vantage point that could clarify the mechanism of protein aggregation in neurodegenerative disease^70,71^ as well as other biological phase separation.

## Acknowledgements

Many thanks to Mariane Le Fur for technical guidance and Eric Gale and Alykhan Shamji for thoughts on the manuscript.

## Funding

This work was supported by a grant from the National Institutes of Health (R01CA241179) and utilized equipment funded by the Massachusetts Life Science Center.

## Competing interests

No conflicts of interest are declared.

## Methods

### Dimer model calculations

All calculations were performed in MATLAB. MATLAB code is available at https://github.com/dubachLab/phaseSeparation. Plots were generated in Graphpad Prism.

### Condensate formation

Unless otherwise noted all chemicals were obtained from Sigma Aldrich. FFome was obtained from Ambeed. Peptides were obtained from Alamanda Polymers or Genscript (4mer peptides). DNA (ACGTATCCGAATGCCAGTTG) and RNA (ACGUAUCCGAAUGCCAGUUG) were obtained from IDT. FFome was dissolved in 5 mM HEPES buffer at pH<4 and 500 mM KCl. pH was controlled by addition of HCl and NaOH or KOH. All peptides and oligonucleotides were reconstituted in 10 mM TRIS buffer, pH 8.0. pH was adjusted using NH_3_OH or HCl. Phase separation was achieved through mixtures of stock solutions to generate the desired concentrations and imaged in 96 well glass bottom well plates (CellVis) on a Nikon Ti2 widefield microscope in differential interference contrast (DIC) with a 60x NA 1.49 oil immersion or 20x NA 0.75 objective.

### Data and materials availability

All data are available in the main text or the supplementary figures. All materials are commercially available.

## Supplementary Information

### Thermodynamic Calculations

Spontaneous dimer binding of any molecules arises through a reduction in free energy with an affinity that is defined by a dissociation constant. The steady-state fraction bound of interacting molecules is a function of the total concentration and the dissociation constant^72^.

For a homo-binding (dimer) interaction, the dissociation constant is defined as:

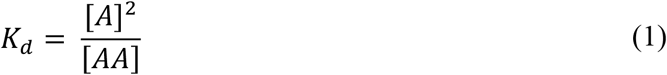

where, [*A*] is the concentration of the free protein *A* and [*AA*] is the concentration of the bound (dimerized) protein. From the total concentration [*A*]_*total*_ (free and bound) in the system:

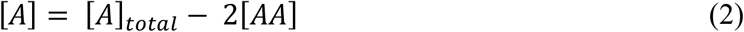

and, through substitution:

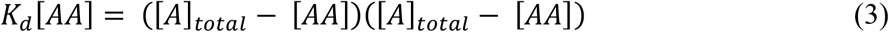

Through the quadratic equation the total bound concentration as a function of total protein and dissociation constant of dimer binding generates the following equation:

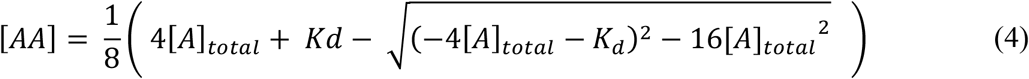

The total free concentration can then be calculated through mass balance.

Liquid phase separation in a system generates both dilute and dense regions with concentrations that are defined by the partition coefficient and the dense phase volume fraction. The partition coefficient is defined as:

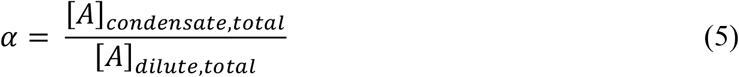

where, *α* is the partition coefficient, [*A*]_*condensate,total*_ is the total concentration of *A* (bound and unbound) in the condensate (dense phase) and [*A*]_*dilute,total*_ is the total concentration of *A* (bound and unbound) in the dilute phase. The concentration in each phase can be calculated from the total concentration in the system. First:

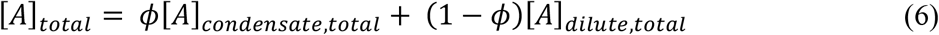

where, *ϕ* is the volume fraction of the dense phase and [*A*]_*total*_ is the total concentration (bound and unbound) in the system. Here, the number and size of condensates are not considered, rather the total volume of each phase. Rearranging, the concentration in the condensate can be calculated from:

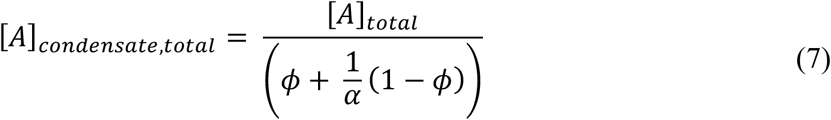

Conversely the dilute phase concentration is:

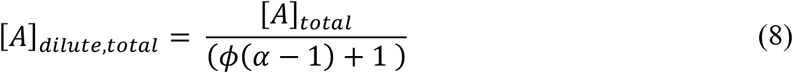

The total bound and unbound concentrations in each phase of the separation (condensate or dilute) can be directly calculated from the total protein concentration within that phase ([*A*]_*condensate,total*_ and [*A*]_*dilute,total*_) and the binding dissociation constant using Equation 4 and mass balance.

By definition, condensates have a higher concentration than diffuse distribution in the absence of phase separation, and thus a higher fraction bound within the condensate, while dilute regions have a lower concentration than the diffuse state, which reduces the fraction bound in the dilute region (**Fig. 2A**).

Summing the volume-weighted, K_d_-governed binding in both regions produces the total fraction bound in the system upon phase separation, which generates an effective dissociation constant:

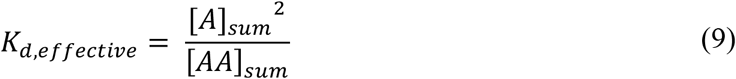

where, [*A*]_*sum*_ and [*AA*]_*sum*_ are the concentrations of unbound and bound protein in the system.

The effective K_d_ is lower in the presence of condensates due to an increase in overall bound protein in the system. In the absence of phase separation, the effective K_d_ is the same as the binding K_d_. The effective K_d_ can then be used to determine the impact of condensation on the Gibbs free energy of the system through the energy of binding:

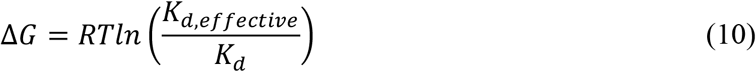

where, Δ*G* is the change in free energy of binding that arises due to the presence of condensates.

However, there is an entropic cost to de-mixing that occurs when condensates are present. The cost is dependent on the concentrations in the dense and dilute regions and can be calculated from:

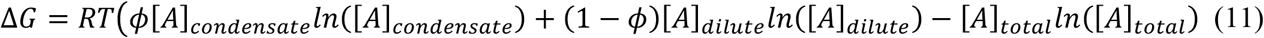

where, [*A*]_*condensate*_ is the concentration of unbound *A* in the condensate, [*A*]_*dilute*_ is the concentration of unbound *A* in the dilute phase, and [*A*]_*total*_ is the concentration of unbound *A* in the absence of condensates. Calculating entropic cost as the change in entropy from the system without condensates also captures the decrease in entropy that arises from a reduction of total species moles that occurs through increased dimer formation. The entropic cost is calculated for each protein species (*A* and *AA*) and summed to obtain the total entropic cost of condensation.

Summing the change in Gibbs free energy of binding and the entropic cost of de-mixing generates the total energetic impact of phase separation. Yet, to be physically realistic a boundary condition that limits the concentration in the dense phase is required. In the absence of a boundary condition, at low affinity and high concentration parameters, energetic minimums occur with dense phase concentrations that are not possible. Therefore, a third thermodynamic force needs to be considered - the chemical potential of the protein. The total chemical potential in the presence and absence of phase separation does not obviously change as the total amount of material is the same. However, chemical potential is a function of density^73^ and is non-ideal, increasing exponentially as the density approaches concentration saturation^74^. Furthermore, the physical size of the protein will limit the achievable concentration in the dense phase. To capture the energetic impact of dense phase concentration dependent chemical potential the following expression is used:

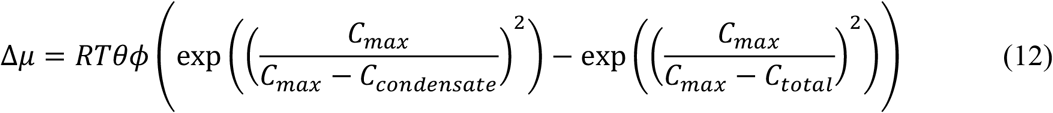

where, Δ*▱* is the change in chemical potential upon condensate formation, *θ* is a constant, C_max_ is the saturation (maximum) total protein concentration in solution that can be achieved, C_condensate_ is the total protein concentration in the dense phase and C_total_ is the total concentration in the system. This representation was used to generate a chemical potential of protein with non-linear dependence on concentration that approaches infinity at the saturation concentration.

Varying the values of *θ* and C_sat_ demonstrates that inclusion of Δ*▱* restricts the condensate concentration to physically achievable values, particularly at higher protein concentrations, but has little impact at lower protein concentrations (Fig. S2 and S3). Thus, C_sat_ and *θ* values were chosen to be 100 mM and 10^−2^, respectively.

For a hetero-binding interaction with a 1:1 stoichiometry, the thermodynamic calculations will be similar except for the mass balance. The dissociation constant is defined as:

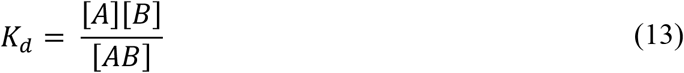

where, [*A*] and [*B*] are the concentrations of free (unbound) proteins *A* and *B*, respectively, and [*AB*] is the bound concentration. The total concentrations are known and thus the concentration of free protein is:

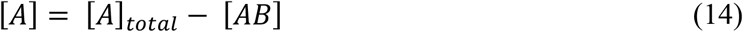

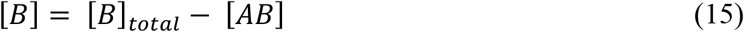

substituting into the previous equation:

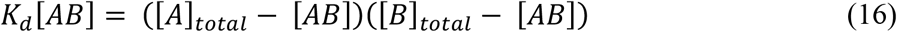

and, through the quadratic formula:

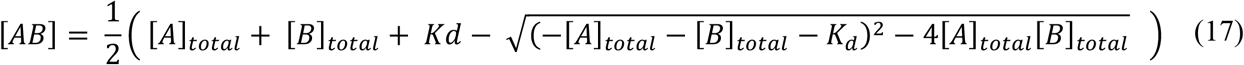

The concentrations of *A* and *B* can subsequently be determined through mass balances. Under phase separation, the concentrations in the dense and dilute phases can be calculated as described from homo-dimerization for both *A* and *B*.

For the calculations here the concentration of *A* and *B* are equal. Thus, the Gibbs free energy of binding in the presence of phase separation can be calculated as described for homo-dimer interactions. The entropic cost of de-mixing is the sum of the costs for *A, B* and *AB*. And the same chemical potential boundary condition is applied.

## Notes

### Competing Interest Statement

The authors have declared no competing interest.

### Summary of Updates

This version provides more supporting results and describes the dimer model from multiple viewpoints to better convey the thermodynamics governing phase separation.

