## Supplementary data for "Dimerization thermodynamics govern biological phase separation"

**Authors:** J. Matthew Dubach

**Affiliations:**
Institute for Innovation in Imaging, Department of Radiology, Massachusetts General Hospital, Harvard Medical School, Boston, MA

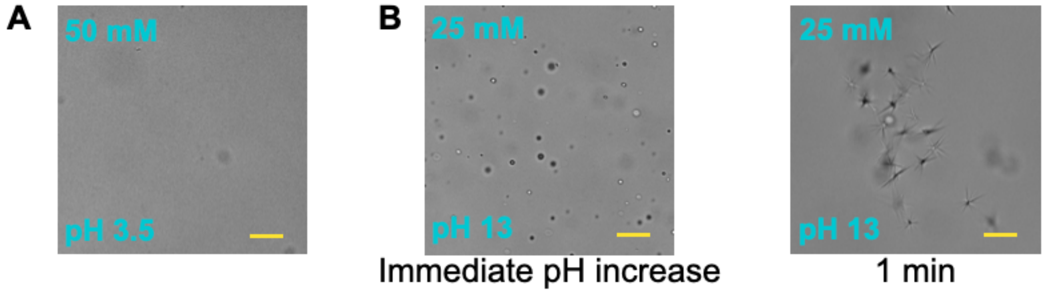


**Supplementary Figure 1. FFome phase separation**. (**A**) Representative DIC image of 50 mM FFome at pH 3.5. (**B**) Representative DIC images of 25 mM FFome at pH 13 immediately after diluting the solution in KOH to raise the pH (left) and after 1 minute (right). Scale bare = 30 μm.

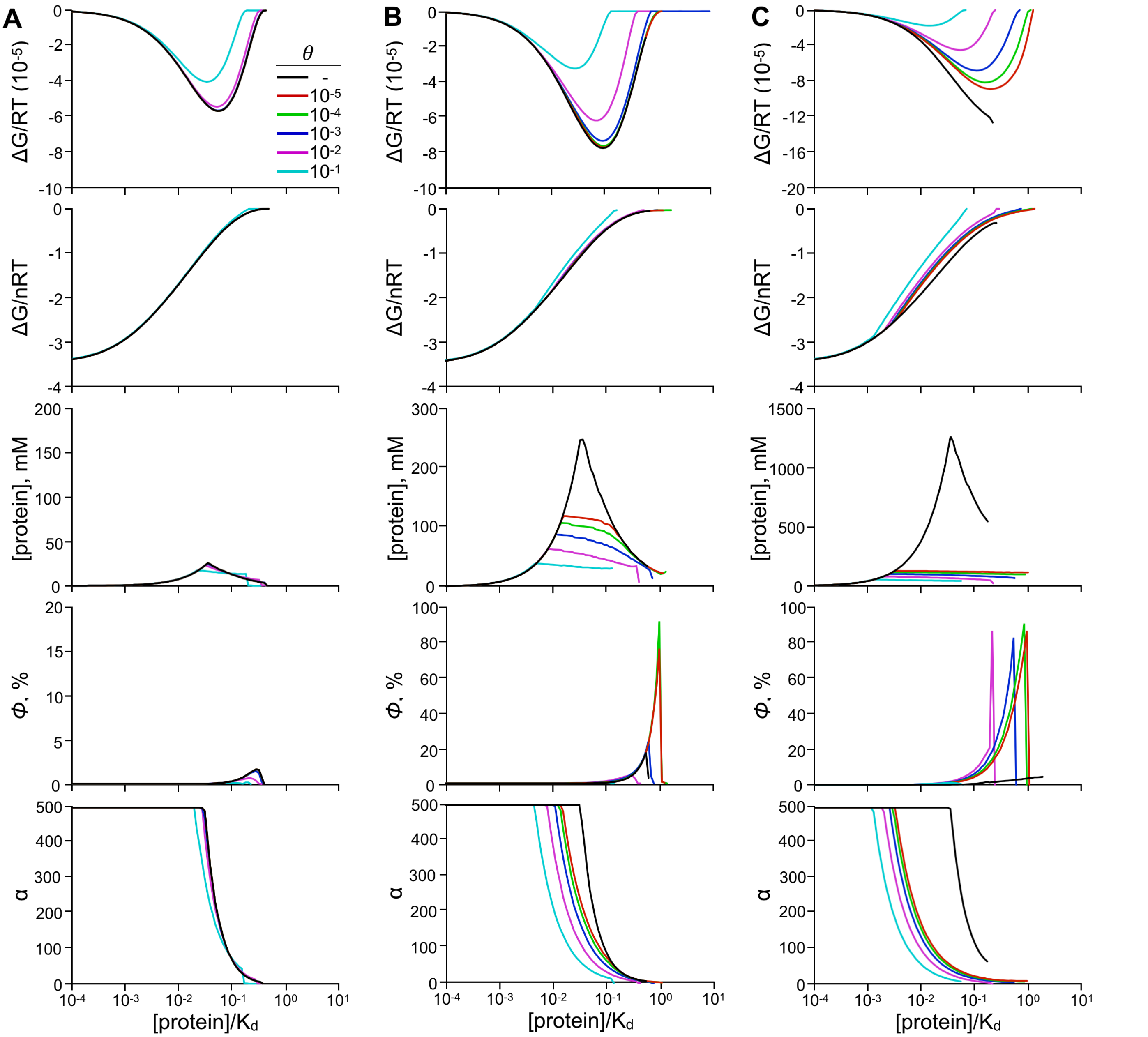


**Supplementary Figure 2. Impact of the non-ideal chemical potential constant on the binding model calculations.** Calculations of minimum total energy, minimum energy per mole, total condensate protein concentration at the energetic minimum, volume fraction at the energetic minimum, and partition coefficient at the energetic minimum as a function of [protein]/$K_d$ (single protein concentration) at multiple constant values for equimolar binding proteins with a $K_d$ of **(A)** 1 mM, **(B)** 10 mM, and **(C)** 50 mM – $C_{max}$ = 200 mM.

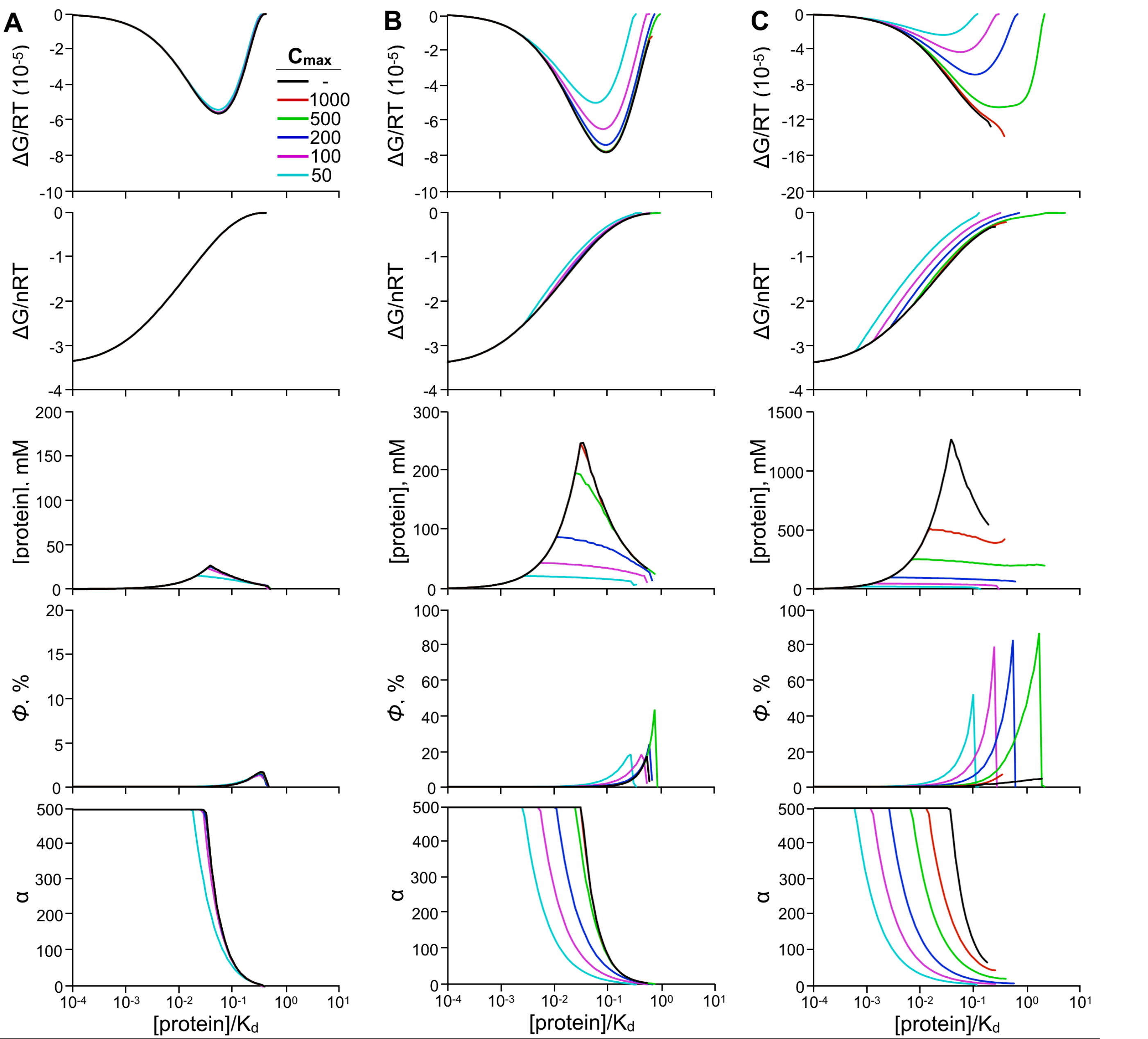


**Supplementary Figure 3. Impact of the non-ideal chemical potential maximum concentration on the binding model calculations.** Calculations of minimum total energy, minimum energy per mole, total condensate protein concentration at the energetic minimum, volume fraction at the energetic minimum, and partition coefficient at the energetic minimum as a function of [protein]/$K_d$ (single protein concentration) at multiple maximum concentration values for equimolar binding proteins with a $K_d$ of **(A)** 1 mM, **(B)** 10 mM, and **(C)** 50 mM – $\theta = 10^{-3}$.

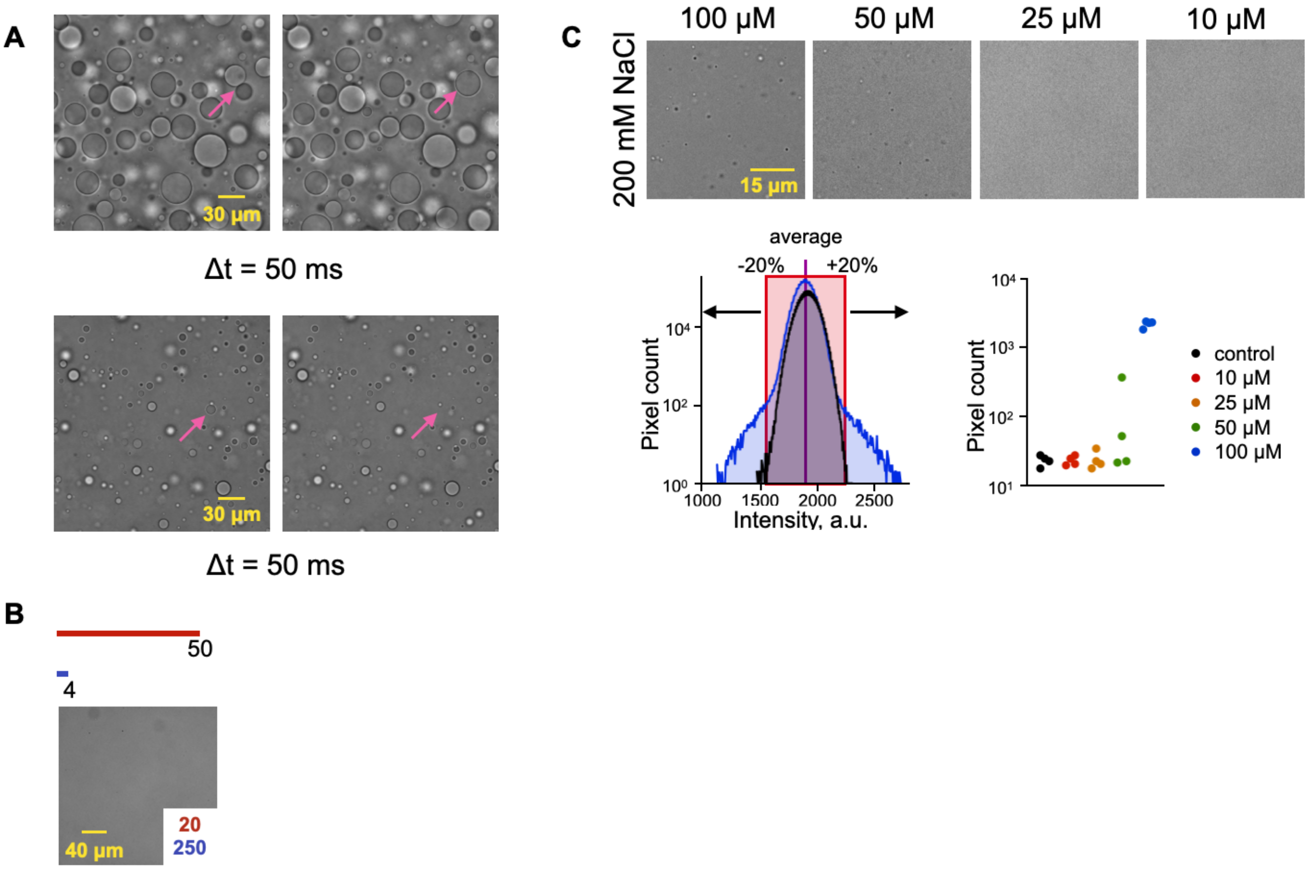


**Supplementary Figure 4. Confirming and identifying condensates**. (**A**) Representative DIC images of 10 mM equimolar $K_{20}$-$E_{20}$ condensates in 200 mM NaCl merging (top) and a 2 mM equimolar $K_{20}$-$E_{20}$ condensate in 40 mM NaCl spontaneously reentering solution, pink arrows identify events – step between frames is 50 ms. (**B**) Representative DIC images $K_{50}$ (red) and $E_4$ (blue) mixtures at equal charge ratios. (**C**) (top) Representative DIC images of equimolar $K_{20}$-$E_{20}$ condensates in 200 mM total NaCl at different equimolar peptide concentrations. (bottom) Representative pixel intensity histograms of 100 µM equimolar $K_{20}$-$E_{20}$ condensates (blue) and solution control (black) images. The pixel count with values outside the average ± 20% was integrated for each image and significant differences (n=4, Student’s t test) from control were determined to identify the presence of condensates in low concentration measurements.

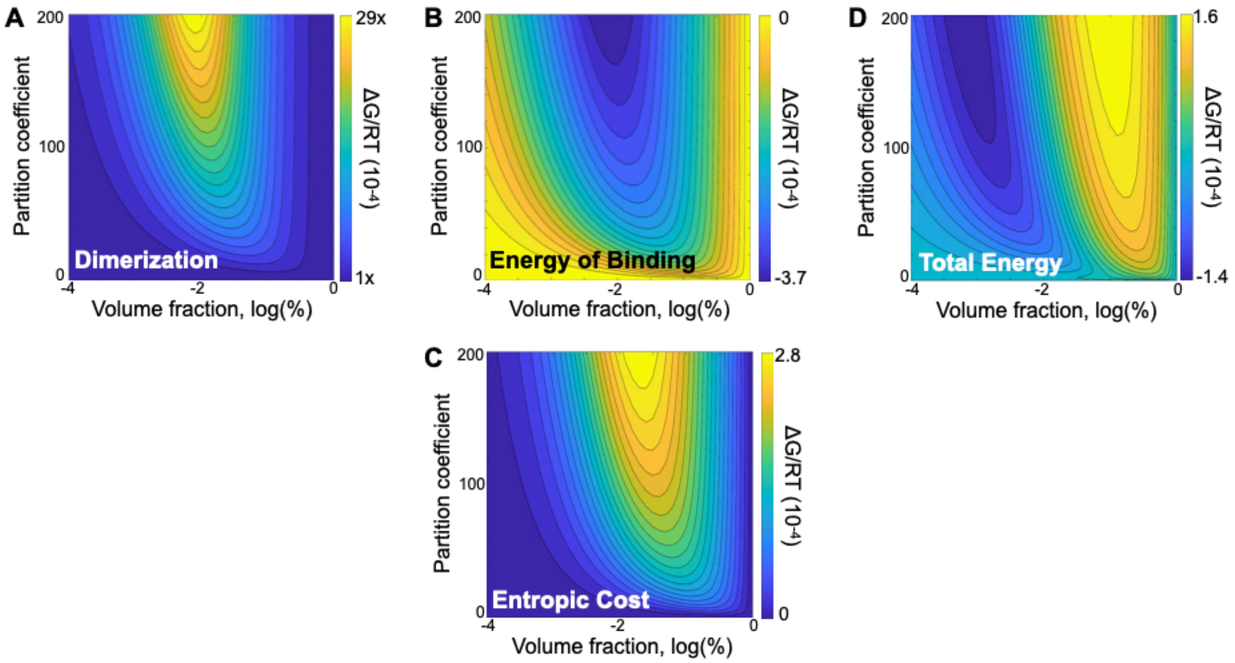


**Supplementary Figure 5. Dimer model energy calculations for hetero-dimer binding.** (**A**) Calculations for total hetero-dimerization fraction bound upon liquid-like phase separation as a function of dense phase volume fraction and partition coefficient. (**B**) Calculations for the change in the Gibbs free energy of binding upon liquid-like phase separation as a function of dense phase volume fraction and partition coefficient. (**C**) Calculations for the change in the entropy of de-mixing upon liquid-like phase separation as a function of dense phase volume fraction and partition coefficient. (**D**) Calculations for the change in total system energy upon liquid-like phase separation as a function of dense phase volume fraction and partition coefficient. For all calculations, $K_d$ = 10 mM.

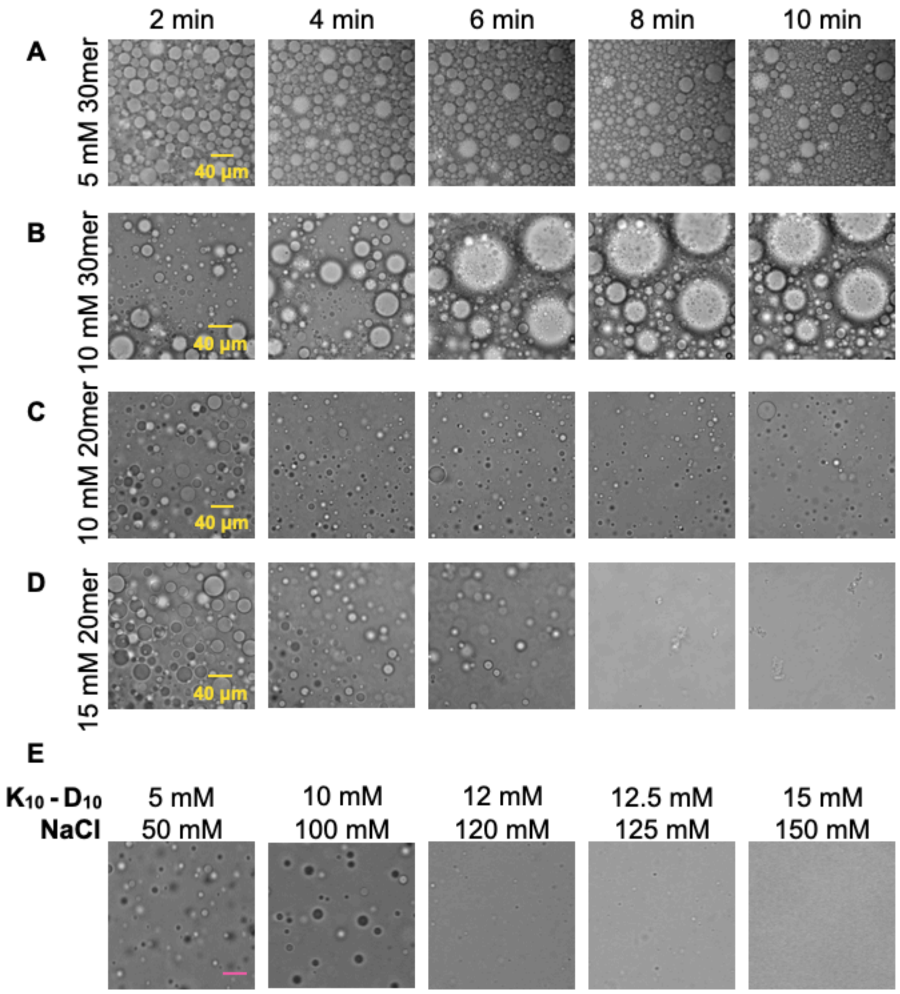


**Supplementary Figure 6. Condensate formation and stability at multiple concentrations.** Representative DIC images of condensates formed by equimolar mixtures of (**A**) 5 mM $K_{30}$-$E_{30}$ at 150 mM NaCl, (**B**) 10 mM $K_{30}$-$E_{30}$ at 300 mM NaCl, (**C**) 10 mM $K_{20}$-$E_{20}$ at 200 mM NaCl, (**D**) 15 mM $K_{20}$-$E_{20}$ at 300 mM NaCl. (**E**) Representative DIC images of condensates formed by equimolar mixtures of $K_{10}$-$D_{10}$, scale bar = 40 µm.

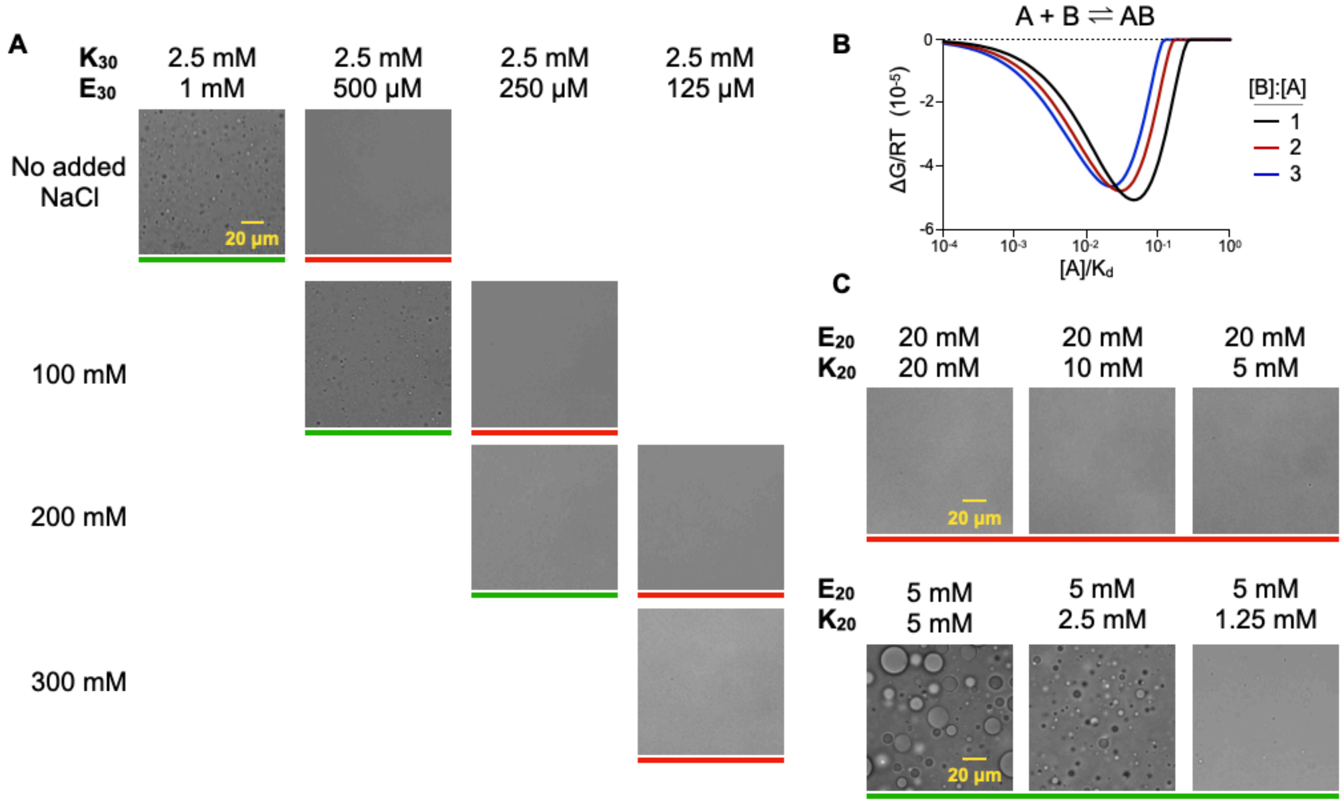


**Supplementary Figure 7. Counterions and concentration govern condensate formation.** (**A**) Representative DIC images of $K_{30}$-$E_{30}$ condensates with no added NaCl (stock counterions only) or added NaCl concentration. Condensates are only present at $K_{30}$-$E_{30}$: 2.5-1 mM no added NaCl, 2.5-0.5 mM 100 mM NaCl, and 2.5-0.25 mM 200 mM NaCl. (**B**) The binding model calculations energy minimum of condensate formation for different ratio mixtures as a function of $[A]/K_d$. Lower ratios of A concentration predict that condensates are no longer energetically favorable at decreasing concentrations of A. (**C**) Representative DIC images of $E_{20}$-$K_{20}$ mixtures. For imbalanced mole ratios an additional 200 mM NaCl was added to the mixture.

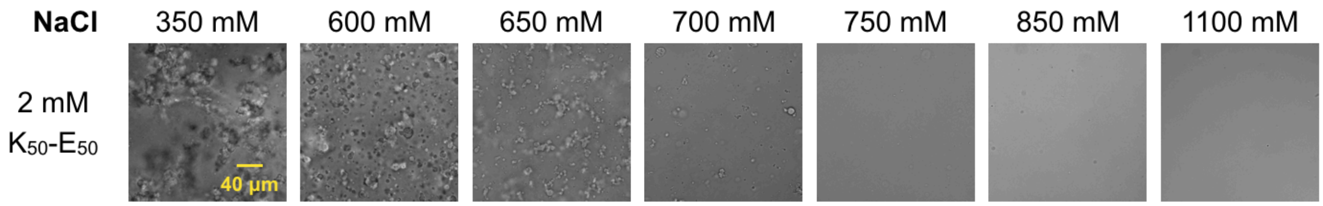


**Supplementary Figure 8. 50mer peptides do not form condensates.** Representative DIC images of 2 mM equimolar $K_{50}$-$E_{50}$ mixtures at multiple NaCl concentrations.

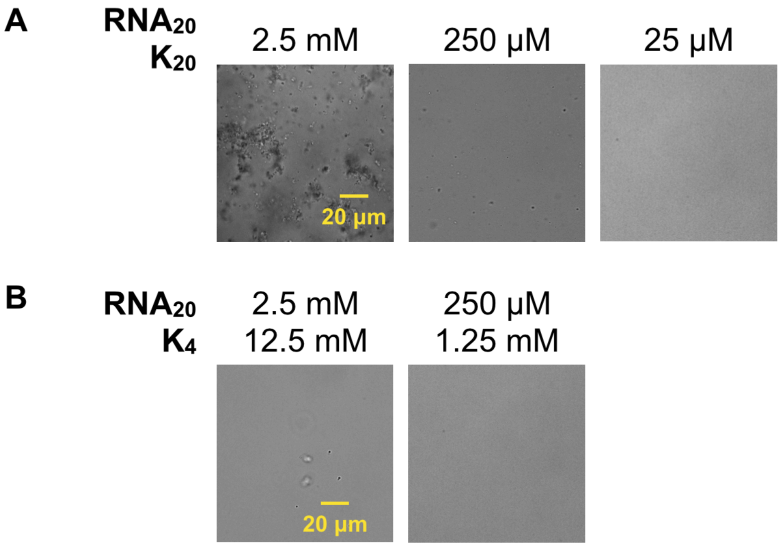


**Supplementary Figure 9. PolyK and $RNA_{20}$ do not form condensates.** (**A**) Representative DIC images of equimolar $K_{20}$ and $RNA_{20}$ mixtures at multiple concentrations. (**B**) Representative DIC images of equal charge mixtures of $K_4$ and $RNA_{20}$.

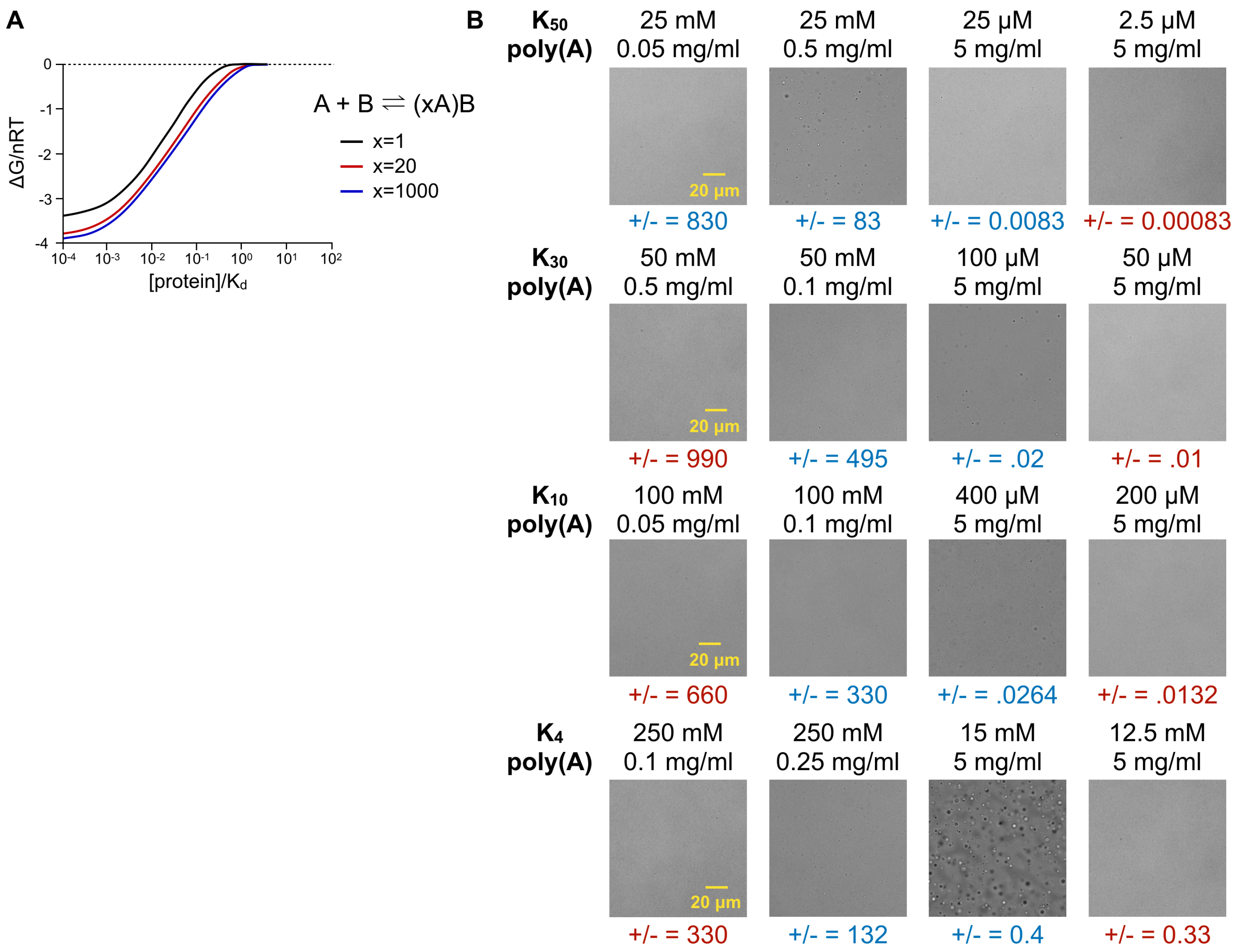


**Supplementary Figure 10. Multiligand polyK binding to poly(A) forms condensates.** (**A**) The binding model calculations of energy minimum per mole for binding interactions where multiple molecules of A can bind B at equimolar mixtures as a function of [Protein]/$K_d$ (single component concentration). (**B**) Representative DIC images of polyK and poly(A) mixtures at multiple mass ratios. Shown are the charge ratio (+/-) and the presence (blue) or absence (red) of condensates.
